# Syd/JIP3 Controls Tissue Size by Regulating Diap1 Protein Turnover Downstream of Yorkie/YAP

**DOI:** 10.1101/2020.04.19.049023

**Authors:** Vakil Ahmad, Gangadhar P. Vadla, Chiswili Y. Chabu

**Author notes:** Corresponding author: C.Y. Chabu.

## Abstract

How organisms control organ size is not fully understood. We found that Syd/JIP3 is required for proper wing size in *Drosophila*. *JIP3* mutations are associated with organ size defects in mammals. The underlying mechanisms are not well understood. We discovered that Syd/JIP3 inhibition results in a downregulation of the inhibitor of apoptosis protein1 (Diap1) in the *Drosophila* wing. Correspondingly, Syd/JIP3 deficient tissues exhibit ectopic cell death and yield smaller wings. Syd/JIP3 inhibition generated similar effects in mammalian cells, indicating a conserved mechanism. We found that Yorkie/YAP stimulates Syd/JIP3 in *Drosophila* and mammalian cells. Notably, Syd/JIP3 is required for the full effect of Yorkie-mediated tissue growth. Thus Syd/JIP3 regulation of Diap1 functions downstream of Yorkie/YAP to control growth.

This study provides mechanistic insights into the recent and perplexing link between *JIP3* mutations and organ size defects in mammals, including in humans where de novo *JIP3* variants are associated with microcephaly.

**Highlights:**

- Syd/JIP3 is required for proper *Drosophila* wing size
- Syd/JIP3 stabilizes Diap1 to inhibit cell death in *Drosophila* and in mammalian cells
- Activation of Yorkie/YAP stimulates Syd/JIP3
- Yorkie-mediated tissue growth is highly sensitive to Syd/JIP3 dosage

## Introduction

The molecular basis of organ size regulation is a central question in developmental biology. Systemic signals such as insulin and growth factors play a key role in determining body and organ size. In *Drosophila*, starvation or genetic mutations in nutrient-sensing genes produce animals with smaller organs via well-defined and evolutionarily conserved mechanisms (Katsuyama et al., 2015; Stocker and Hafen, 2000). In addition to these systemic factors, transplantation experiments in salamander and in *Drosophila* have revealed that tissues possess intrinsic size control mechanisms. Salamander limbs maintain their size when transplanted onto animals with different limb sizes (Twitty V. C., 1931). Similarly, fragments of larval wing discs or young intact wings grow to reach normal size when transplanted into the abdomen of adult flies (Bryant and Levinson, 1985). In contrast to systemic signals, these tissue-intrinsic organ size control mechanisms are not well understood.

The *Drosophila* wing imaginal disc is a powerful experimental system to understand growth control mechanisms. Embryonically derived cells undergo several rounds of cell proliferation to form a mature epithelium that ultimately gives rise to an adult wing with stereotyped size and shape (Day and Lawrence, 2000; Fristrom and Chihara, 1978; Garcia-Bellido, 1975; Garcia-Bellido and Merriam, 1971; Mandaravally Madhavan and Schneiderman, 1977; Martin et al., 2009; Milan et al., 1996; Worley et al., 2013). Genetic screens aimed at identifying mutations that deregulate growth in this system have led to the discovery of key and broadly conserved tissue-intrinsic growth control signaling pathways, including the Hippo pathway (reviewed in (Hariharan, 2015; Irvine and Harvey, 2015; Pan, 2007). In fly and mammalian tissues, Hippo signaling potently controls tissue size by modulating the activity of Yorkie/YAP, a transcriptional co-activator for cell proliferation and cell survival genes (Dong et al., 2007; Huang et al., 2005; Jia et al., 2003; Oh and Irvine, 2008; Pan, 2007). An earlier study reported that the c-Jun N-terminal kinase (JNK, also known as Bsk–FlyBase) regulates Hippo in the developing wing to control wing size. Localized JNK activity represses Hippo to allow Yorkie/YAP to drive cell proliferation throughout the tissue, hence promoting wing growth (Willsey et al., 2016). The Hippo-Yorkie/YAP signal is a central tissue-size regulator, and its function is well conserved across species. While research on Yorkie/YAP have generated significant mechanistic insights into how Yorkie/YAP and its transcriptional program are regulated, much less is known about developmental mechanisms for restraining Yorkie/YAP-mediated tissue growth to achieve organ size stereotypy(Bernhard et al., 2000; Pan, 2007; Zheng and Pan, 2019).

Mitogen activating protein kinase (MAPK) scaffolding proteins mediate cellular responses to apoptotic or proliferative signals. Sunday Driver (Syd) or JNK-interacting protein-3 (JIP3) belongs to a family of conserved scaffolding proteins that bind to and facilitate JNK activation (Davis and Tapon, 2019; Davis, 2000; Kelkar et al., 2000). Syd/JIP3 also regulates vesicle transport in worm and mammalian cells (Caswell and Dickens, 2018; Choudhary et al., 2017; Edwards et al., 2013; Gowrishankar et al., 2017; Sun et al., 2017; Taylor and Alessi, 2020).

Using the developing *Drosophila* wing as a model system, we found that Syd/JIP3 is required for proper wing size. Mechanistic studies revealed that Syd/JIP3 functions downstream of Yorkie/YAP by controlling the stability of Diap1, an important cell survival regulator and Yorkie/YAP target. Syd/JIP3 regulates Diap1 post-transcriptionally by controlling its protein turnover. Congruent with these observations, knocking down Syd/JIP3 has no effect on Diap1 transcription but leads to a loss of Diap1 protein and correspondingly causes ectopic cell death, resulting in a smaller wing. Restoring Diap1 protein in Syd/JIP3 knockdown tissues restores wing size. Importantly, activation of Yorkie/YAP upregulates Syd/JIP3 protein and interfering with this stimulation suppresses Yorkie/YAP-mediated tissue overgrowth. In addition to defining a novel growth control mechanism, this work provides molecular insights into the perplexing association between *JIP3* mutations and small brains in mice and humans (Kelkar et al., 2003; Platzer et al., 2019).

## Materials and Methods

### Fly strains

All the crosses were maintained at 25°C unless otherwise described. Fly strains used for this study are: *UAS-Syd-RNAi* (Bloomington Stock #43232), *UAS-Syd-RNAi* (VDRC Stock #35346), *UAS-dSyd.GFP* (Mary K. Baylies), *Df(3L)syd*^*A2*^ (Bloomington Stock #32017), *syd*^*PL00426*^ (Bloomington Stock #19495), *UAS-Bsk-DN* (Tian Xu), *UAS-Yki* (Doujia Pan), *UAS-Yki*^*S111A.S168A.S250A*^ (Bloomington Stock #28817), *UAS-Hpo-RNAi* (Bloomington Stock #33614), *UAS-Diap1* (Bloomington Stock #6657), *diap1-LacZ* (Bloomington Stock #12093), *diap1^33-1s^*(Hermann Steller), *fj-LacZ* (Bloomington Stock #44253), *CycE-lacZ* (Bloomington Stock #59065), *ex-lacZ* (Bloomington Stock #44248), *Jub-GFP* (Bloomington Stock #56806), *dronc^51^*(Bloomington Stock #23284) including GAL4 drivers *Dcr2,engrailed-Gal4, nubbin-Gal4, patched-Gal4, hedgehog-Gal4 and gmr-Gal4.*

### Immunohistochemistry

Wing imaginal discs were dissected, fixed, and stained as previously described (Chabu and Xu, 2014; Pagliarini and Xu, 2003). The primary antibodies used were rabbit anti-PH3 (1:1000, Sigma); rabbit anti-Diap1 (1:1000, Hermann Steller); rabbit anti-pWts (1:200, D. Pan), mouse anti-β-gal (1:1000, Sigma). Secondary antibodies were from Invitrogen. Specimens were imaged on a Leica TCS SP8 confocal microscope. Images were analyzed and processed with IMARIS (Bitplane, Switzerland) and illustrator software (Adobe).

### Western blots

Wing imaginal discs from third-instar larvae were dissected and homogenized in lysis buffer (50 mM HEPES pH 7.5, 150 mM KCl, 5 mM MgCl_2_, 0.01% Triton-X) supplemented with protease and phosphatase inhibitor cocktail (Cat #78443, Thermo Scientific). For cell culture, protein lysates were prepared in lysis buffer (20 mM Tris-HCl pH-7.5, 150 mM NaCl, 1 mM Na2EDTA, 1 mM EGTA, 1% Triton, 2.5 mM sodium pyrophosphate, 1 mM β-glycerophosphate, 1 mM Na3VO4, 1 μg/ml leupeptin) supplemented with protease and a phosphatase inhibitor cocktail (Cell Signaling #9803S). Proteins were electrophoresed on SDS-PAGE using 4-15% Mini-PROTEAN^®^ TGX™ precast gel (Cat #456-1085, Bio-Rad), transferred onto Nitrocellulose membranes, and incubated with rabbit anti-Diap1 (1:1000, Hermann Steller) or mouse anti-JIP3 (1:1000, Cat #46663, Santa Cruz), mouse anti-HA (1:1000, Cat #7392, Santa Cruz), mouse anti-c-IAP(1:1000, Cat #271419, Santa Cruz), rabbit anti-β-Actin (1:1000, Cat #4967, Cell Signaling Technology), mouse anti-GAPDH (0.3 μg/ml, Cat #2G7, DSHB). Secondary antibodies used are anti-mouse horseradish peroxidase (HRP) (1:5000, Cat #31430, Invitrogen) and anti-rabbit HRP (1:10000, Cat #31460, Invitrogen). Antibody signals were detected using the ECL kit (Cat #32106, Thermo Fisher scientific) and ChemiDoc− MP Imaging System (Bio-Rad).

### Adult wing imaging and measurements

Adult wings of female animals were dissected, mounted, flattened with glass slide, and imaged at 1.0X or 2.5X using Leica Microsystems, DMC 4500. In supplementary Figure 1, female flies were washed with 75% ethanol and then with 95% ethanol for 2 min. Animals were dried on blotting paper in oven at 50°C for 2 min. Wings were removed and mounted on a glass slide in mineral oil, covered with 22 × 40 mm coverslip and imaged at 2.5X. Areas of the adult wings were measured using the *freehand selection* and *analyze* tools of ImageJ software 1.50i. P-values were calculated using the two-tailed t-test function in Microsoft Excel.

### Cell death detection in *Drosophila*

For labeling dead cells with TUNEL assay, ApopTag® Red In Situ Apoptosis Detection Kit (Cat # S7165) was used according to manufacturer’s instruction. Briefly, dissected wing discs were fixed and incubated in equilibration buffer at room temperature for 20 min followed by incubation with terminal deoxynucleotidy transferase in reaction buffer for 1 h at 37°C. Reaction was stopped using stop buffer diluted in dH_2_O. Disc were incubated with conjugated anti-digoxigenin antibodies for 2 h at room temperature, washed, and mounted using Vectashield mounting medium.

### Cell lines and cell culture

Authenticated Human Embryonic Kidney 293 (HEK-293) cells were cultured in Dulbecco’s Modified Eagle Medium (#11965-092) supplemented with 10% FBS. TrypLE (Gibco#12604-021) was used for immobilizing cells. Cells were counted in a Neubaur cell counter chamber following Trypan Blue dead cell exclusion (Cat #T8154, Sigma). Dharmacon on-target siRNA-SMARTpool plus Human scramble control (Cat #D-001810-10-05, Dharmacon) or JIP3 (Cat #L-003596-00-0005, Dharmacon) were used for RNA interference experiments in HEK-293 cells. HEK 293 cells were transfected with purified pCI-HA-YAP plasmid (Cat #27007, Addgene). DharmaFECT (Cat #T2002-02, Dharmacon) was used for transfection. Protein knockdown was assessed 72 hours after transfection.

### Real-time RT-PCR

*Drosophila* wing tissues were dissected from 72-96 hours after egg-laying (AEL) or from 120-150 hours AEL wild-type animals for Syd/JIP3 quantitative real-time PCR (qRT-PCR). EZ Tissue/Cell culture total RNA Miniprep kit (Cat #R1002-50, Bio Research) was used to extract RNA in biological triplicates. The cDNA Reverse Transcription kit (Cat #4368814, Applied Biosystem) was used to generate cDNA from 100 ng total RNA. SYBR Green iTaq Supermix (Bio-Rad) was used for qRT-PCR. Expression of housekeeping gene *Rp49* (RpL32–FlyBase) was used as an internal control. Primer sequences are as follows:

*Syd*: 5’-ACTTGACTCTACGCTGCGGCTGT-3’ and 5’-GGTGCCGATCCACAGCCGGT-3’

*rp49*: 5′-GGCCCAAGATCGTGAAGAAG-3′ and 5′-ATTTG-TGCGACAGCTTAGCATATC-3′

### Flow cytometry analysis

Flow cytometric detection of the cell death dye propidium iodide (PI) was used. PI was used at 10 μg/ml in PBS containing 1% BSA. PI was detected at 488/636 nm (excitation/emission) on an ADP Beckman Coulter flow cytometer.

### Statistical analysis

All *in vitro* experiments were reproducibly performed using biological triplicates. A student’s *t-test* was used for comparing the difference between two groups, and a *p* < 0.05 was considered as statistically significant.

## Results

We knocked down Syd/JIP3 in the developing wing (*nubGAL4>Syd-RNAi*) and this resulted in smaller adult wings compared to controls (Figure 1A-E). A second Syd-RNAi line (v#35346) similarly suppressed wing size (Supplemental Figure 1). Consistent with Syd/JIP3 inhibition, larval wing tissues expressing either *Syd-RNAi* construct showed a reduction of Syd/JIP3 protein in Western blot analyses. Anti-Syd/JIP3 Western blots detected two bands: a prominent ~147KD band and a much weaker 135KD band in lysates derived from wild-type wing tissues. These bands were considerably reduced in lysates obtained from wing tissues expressing *Syd-RNAi* (Figure 1F Supplemental Figure 8A, C.). In addition, heterozygosity of two independent null alleles of *Syd* (Bowman et al., 2000; Hacker et al., 2003) (*syd^PL00426/^*^A2^) also yielded animals with smaller wings compared to controls (Figure 2F, K). Taken together these data indicate that Syd/JIP3 regulates wing size.

**Figure 1.**
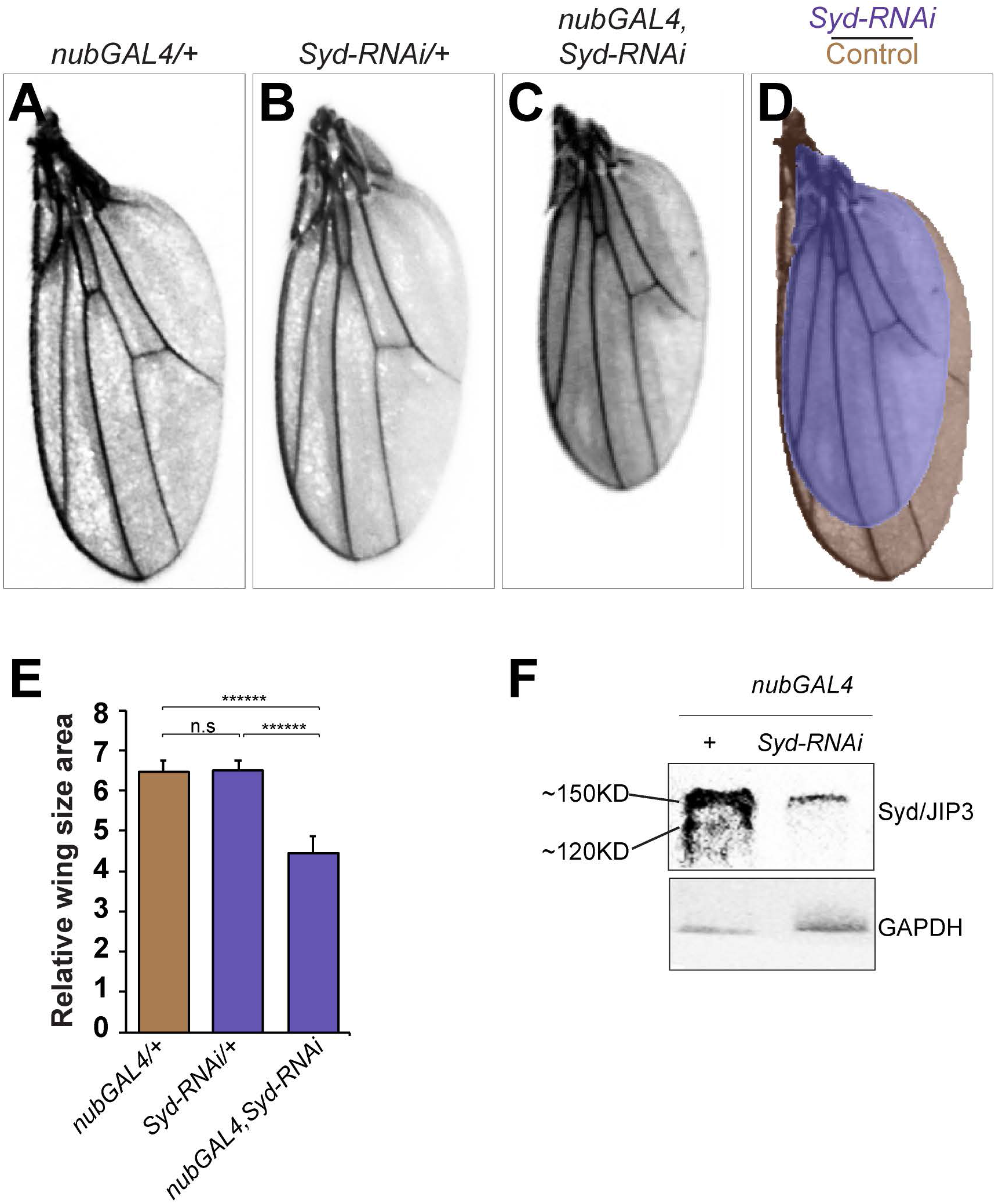
Syd/JIP3 regulates adult wing size. (A-D) Representative images showing adult wings from female animals of the indicated genotypes. Wings from *nubGAL4* or *Syd-RNAi* parental lines are shown in (A) and (B), respectively. (C) Image showing the effect of Syd knockdown (*nubGAL4*, *Syd-RNAi*). (D) Image of a pseudo-color overlay of (A) (shown in brown) and (C) (shown in light indigo). (E) Quantification of wing size. For this and all subsequent quantification figures, plotted values indicate the mean ±SD. The number of tissues analyzed (N) per genotype is indicated inside the bar. All experiments were repeated at least twice. *P*-value is computed from a t-test: p-value < 10^−20^ = ******, p-value = 10^−20^– 10^−10^ = *****, p-value = 10^−10^ –10^−5^ = ****, p-value = 10^−5^ – 10^−3^ = ***, p-value=10^−3^ – 10^−2^ = **, p-value = 0.01 – 0.05 = *, p-value ≥ 0.05 = not significant, ns. (F) Western blot image showing Syd/JIP3 protein abundance in third-instar wing disc lysates derived from animals expressing *RFP* alone (control) or co-expressing *Syd-RNAi* in the whole disc proper using *nubGAL4*. Blots were stained with anti-Syd/JIP3 or GAPDH (loading control) antibodies.

**Figure 2.**
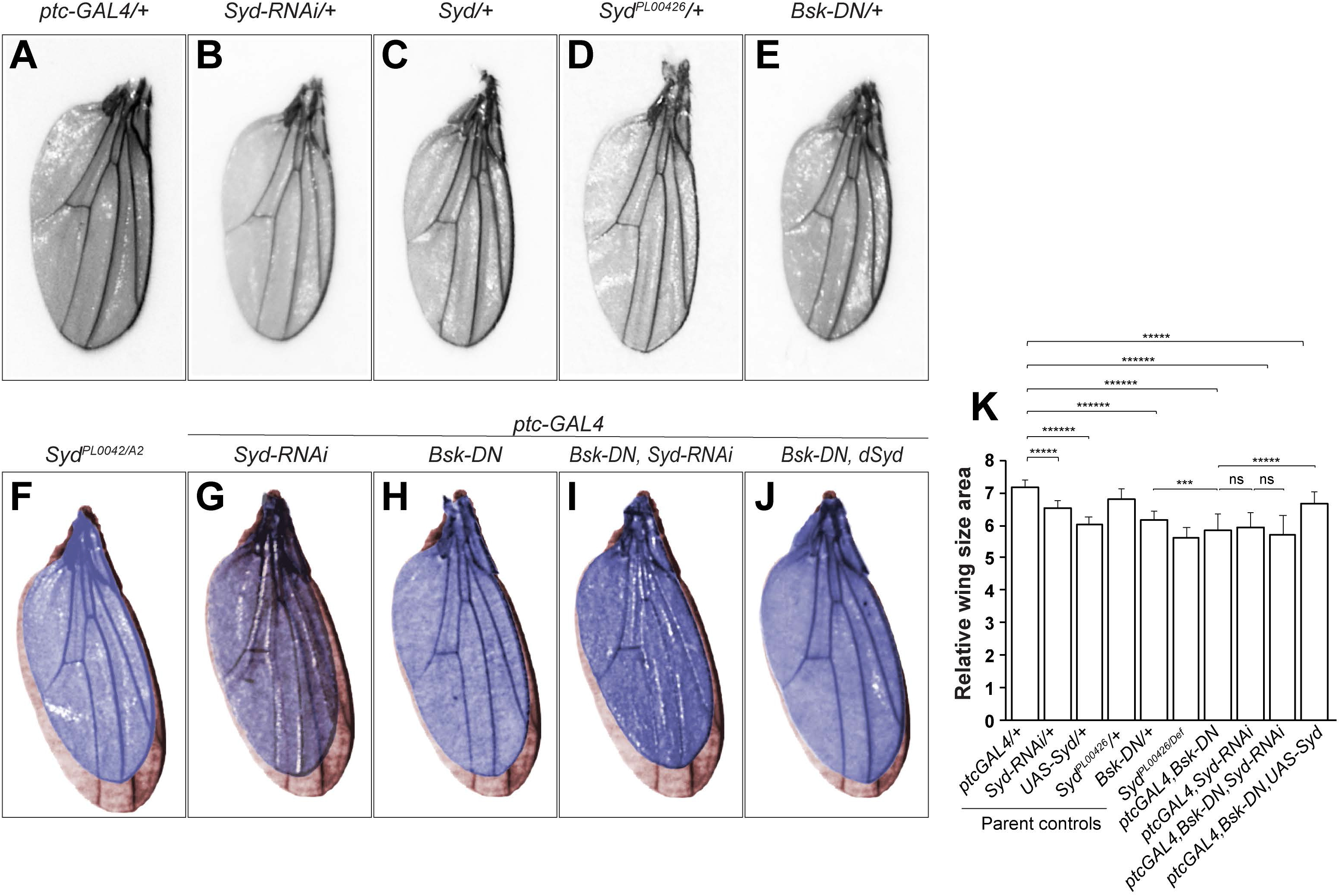
Syd/JIP3 functions downstream of JNK. (A-J) Representative images showing adult wings from female animals of the indicated genotypes. Wing images from control parental fly lines are shown in (A-E). (F-J) Pseudo-color overlay of wing images dissected from *ptcGAL4* control (brown) or from *Syd*^*PL00426*^/*Syd*^*A2*^ trans-heterozygote animals (indigo) (F) or animals expressing *Syd-RNAi* (G) or *Bsk-DN* (H) or co-expressing *Syd-RNAi* and *Bsk-DN* (I) or wild-type *Syd* and *Bsk-DN* (J) all under the control of *ptcGAL4*. (K) Quantification of (A-J).

Interestingly, *JIP3* loss of function mutations are associated with brain size defects in mice and humans (Kelkar et al., 2003; Platzer et al., 2019). We sought to determine the mechanism by which Syd/JIP3 regulates tissue size.

The wing disc consists of a posterior (P) and an anterior (A) compartment. The patterning gene *engrailed* is specifically expressed in the P compartment of the wing disc where it induces Hedgehog (Hh) expression, while transcriptionally repressing Hh target genes (Garcia-Bellido, 1975; Tabata et al., 1992; Zecca et al., 1995). Hh protein is transported via cell-to-cell exocytosis-endocytosis cycles to the A compartment, where it activates Cubitus interuptus (Ci or Gli in vertebrates). In turn, Ci stimulates the expression of dTRAF1 (*Drosophila* tumor necrosis factor receptor-associated factor-1) a JNK pathway component, leading to JNK signaling activation at the P/A boundary. This localized JNK activity acts via the LIM domain family protein Ajuba to repress Hippo signaling throughout the entire tissue (Sun and Irvine, 2013; Willsey et al., 2016). It is unclear how this local effect on Hippo signaling is transferred throughout the whole tissues to promote tissue growth.

We asked whether Syd/JIP3 regulates wing size via this Ci-JNK-Hippo signaling cascade. We turned to *ptc-GAL4* to knockdown Syd/JIP3 in P/A boundary cells where JNK is normally active. We noted that *ptc-GAL4* control wings were larger than their *nub-GAL4* counterpart controls (Figure 2A, K versus Figure 1A, E). *ptcGAL4-*driven expression of *Syd-RNAi* generated wings that were smaller than either parental control (*ptcGAL4 or Syd-RNAi*) wings (Figure 2G, K). These findings raised the possibility that Syd/JIP3 regulates wing size via JNK.

To directly test this hypothesis, we performed genetic epistasis experiments between Syd/JIP3 and JNK. We used the P/A boundary driver *ptcGAL4* to express a potent JNK inhibitor (dominant-negative version of JNK, *Bsk-DN*) alone or in the presence of *Syd-RNAi* and determined the effect of these manipulations on adult wing size. As expected, *Bsk-DN* reduced wing size compared to control (Figure 2H, K). This size reduction effect was not as pronounced as what we had observed with *Syd-RNAi* knockdown at the P/A boundary or throughout the wing tissue (*nubGAL4, Syd-RNAi*) (Figure 1A-E and Figure 2G, H, K). Importantly, blocking JNK failed to exacerbate the small wing phenotype caused by Syd knockdown (*ptcGAL4>Bsk-DN, Syd-RNAi*). Instead, these wings resembled Syd-depleted wings (Figure 2G-I, K). Moreover, overexpression of wild-type version of *Syd* rescued the *Bsk-DN* small wing phenotype (Figure 2H, J, K). Collectively, these findings argue that Syd/JIP3 is epistatic to JNK and that Syd/JIP3 functions downstream of JNK.

JNK acts via the LIM domain family protein Ajuba to repress Hippo signaling and promotes wing growth (Willsey et al., 2016). Ajuba represses Hippo by sequestering Warts, a negative regulator of Yorkie/YAP, to cell-cell junctions. This sequestration increases Yorkie/YAP activity, which transcriptionally stimulates CyclinE and the E3-ubiquitin ligase *Drosophila* inhibitor of apoptosis (Diap1) to simultaneously promote cell proliferation and inhibit cell death, hence promoting tissue growth. How this Yorkie/YAP activation propagates throughout the whole tissue remains unclear. Compared to wild-type controls, depleting Ajuba in the developing wing results in smaller adult wings (Das Thakur et al., 2010; Marie et al., 2003; Rauskolb et al., 2014), similar to Syd knockdown. We asked whether Syd is required for the junctional sequestration of Ajuba or Warts and found that it is not. RNAi depletion of Syd in P/A boundary cells (*ptcGal4>Syd-RNAi*) did not reduce Ajuba protein levels or alter its subcellular localization (Supplemental Figure 2A, B, *N=*22 discs). Similarly, the subcellular localization of Warts in Syd knockdown cells was indistinguishable from wild-type (Supplemental Figure 2A, C*, N=22* discs).

Next, we asked whether Syd/JIP3 acts downstream of Yorkie/YAP. *Patched is* expressed in a well-defined 8-10 cells region along the P/A boundary. We used the *ptc-GAL4* promoter to co-express *Yorkie* in this region. The *ptc-GAL4* line harbors a *UAS-GFP* construct in the background, making it possible to quantitatively examine the effect of transgenes expression on tissue growth by scoring the size of the GFP-positive tissue in confocal images. As expected, *ptc-GAL4* driven expression of *Yorkie* resulted enlarged the P/A region compared to controls (Figure 3A, B, I) (Huang et al., 2005). Co-expression of *Syd-RNAi* suppressed Yorkie-mediated tissue overgrowth (Figure 3B, C, I). The suppressive effect of Syd knockdown on Yorkie-mediated tissue overgrowth is not specific to the P/A domain. *Syd-RNAi* also suppressed Yorkie-mediated tissue overgrowth when Yorkie was over-expressed in the posterior compartment or throughout the larval wing pouch (Supplemental Figure 3).

**Figure 3.**
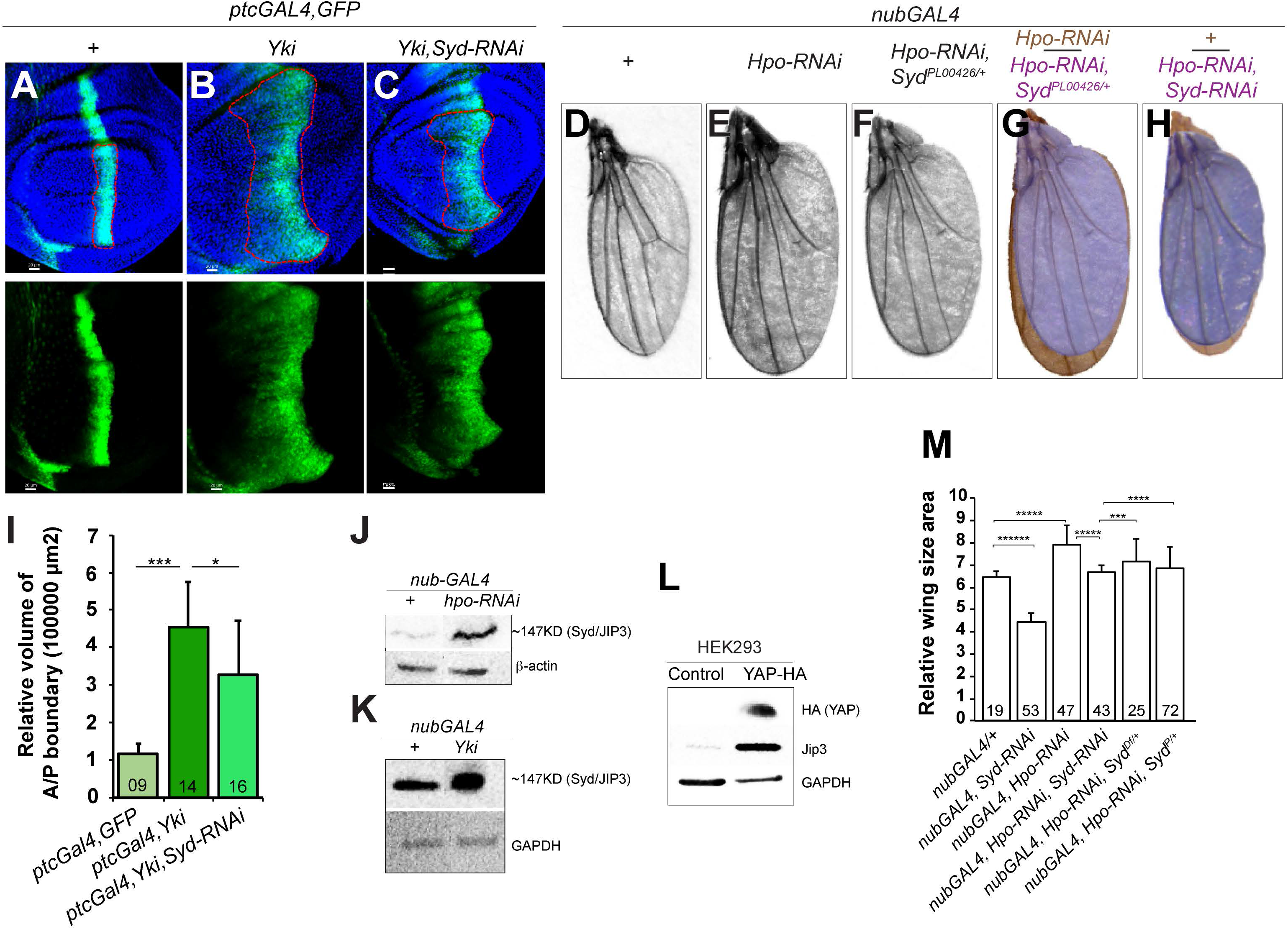
Syd/JIP3 functions in the Hippo pathway. (A-C) Images of wing imaginal discs from age-controlled animals expressing *GFP* alone (A) or co-expressing *Yorkie* (B) or co-expressing *Yorkie* and *Syd-RNAi* (C) under the control of *ptc-GAL4*. Under wild-type condition *ptc-GAL4* expresses GFP in 8-10 cells along the posterior-anterior boundary (A). Overexpression of Yorkie causes expands the size of the GFP-positive region (B). Syd knockdown in Yorkie reduces the Yorkie-mediated tissue overgowth (C). The dotted lines denote the posterior-anterior domain where transgenes were introduced. Size quantification of the GFP-positive tissue across experimental conditions using confocal images and the image data analysis platform IMARIS@BITPLANE data(E). (D-H) Representative images showing adult wings from female animals of the indicated genotypes. Compared to *nubGAL4* control wings (D), expression of *Hippo-RNAi* increases wing size (E). *Syd* haploinsufficiency (*Syd^PL00426/+^*) suppresses the size effect of *Hippo* - *RNAi* expression (F). Pseudo-color overlay wing images are shown in (G and H). (G) Wings expressing *Hippo-RNAi* in the absence or presence of Syd haploinsufficiency (*Syd^PL00426/+^*) are shown in brown and indigo, respectively. (H) Wings expressing *Hippo-RNAi* in the absence or presence of *Syd-RNAi* are shown in brown and indigo, respectively. (I) Quantification of relative volume of *GFP*-positive tissue in A-C. (J) Western blot image showing Syd/JIP3 protein abundance in dissected wing disc lysates from control (*nub-GAL4*) or Hippo-depleted (*nubGAL4>Hpo-RNAi*) tissues. Blots were stained with anti-Syd/JIP3 or anti β-actin (loading control). (K) Western blot image showing Syd/JIP3 protein abundance in dissected wing disc lysates from control tissues (*nub-GAL4*) or tissues over-expressing Yorkie (*nubGAL4>Yorkie*) tissues. Blots were stained with anti-Syd/Jip3 or GAPDH antibodies (loading control). (L) Western blot images showing the effect of YAP overexpression on JIP3 protein abundance in HEK293T cells. HEK 293T cells were transfected either with scramble control or YAP-HA construct. Blots were stained with anti-JIP3, anti-HA (to detect YAP-HA) or or anti-GAPDH (loading control). (M) Wing size quantification of (D-H).

In a complementary approach, we activated Yorkie/YAP by repressing Hippo. In contrast to direct Yorkie/YAP overexpression, this approach activates Yorkie/YAP at physiologically relevant levels. We asked whether knocking down Syd in this context would also repress Yorkie/YAP-mediated tissue overgrowth. As expected, Hippo knockdown resulted in larger wings compared to control (Figure 3D, E, M). *Syd-RNAi* suppressed the large wing effect of Hippo inhibition (Figure 3E, F, H, M). Moreover, heterozygosity of two independent *Syd* mutant alleles (*syd*^*PL00426/+*^ or *syd^A2/+^*) was sufficient to significantly suppress *Hippo-RNAi* large wing phenotype (Figure 3E, G, M).

Collectively, the above findings suggested that Syd/JIP3 regulates wing size downstream or in parallel to Yorkie/YAP. To distinguish between these possibilities, we asked whether Yorkie activation stimulates Syd/JIP3 and it does. Western blot analyses showed that *Hippo-RNAi* triggers the upregulation of Syd/JIP3 protein in *Drosophila* wing tissues (Figure 3J). In addition, expression of Yorkie/YAP elevated Syd/JIP3 protein levels in *Drosophila* and mammalian (HEK293T) cells (Figure 3K, L, and supplemental Figure 8C).

We sought to delineate the underlying mechanism. We examined the expression levels of two key Hippo/Yorkie signaling transcriptional targets, Cyclin E (CycE) and Diap1 using standard transcriptional reporter lines (CycE-LacZ and diap1-LacZ) (Jones et al., 2000; Zhang et al., 2008). Interestingly, Diap1 and CycE expression levels were indistinguishable between wild-type and *Syd-RNAi* expressing cells (Figure 4A, B and Supplemental Figure 4A-C). We extended these analyses to include two additional Hippo signaling transcriptional targets *four-jointed* (fj) and *expanded* (Ex) (Cho et al., 2006; Hamaratoglu et al., 2006) and found that Syd knockdown does not affect the expression of *fj* or *Ex* (Supplemental Figure 4D-I). Thus, it is unlikely that Syd/JIP3 controls wing size by regulating the Hippo transcriptional program.

**Figure 4.**
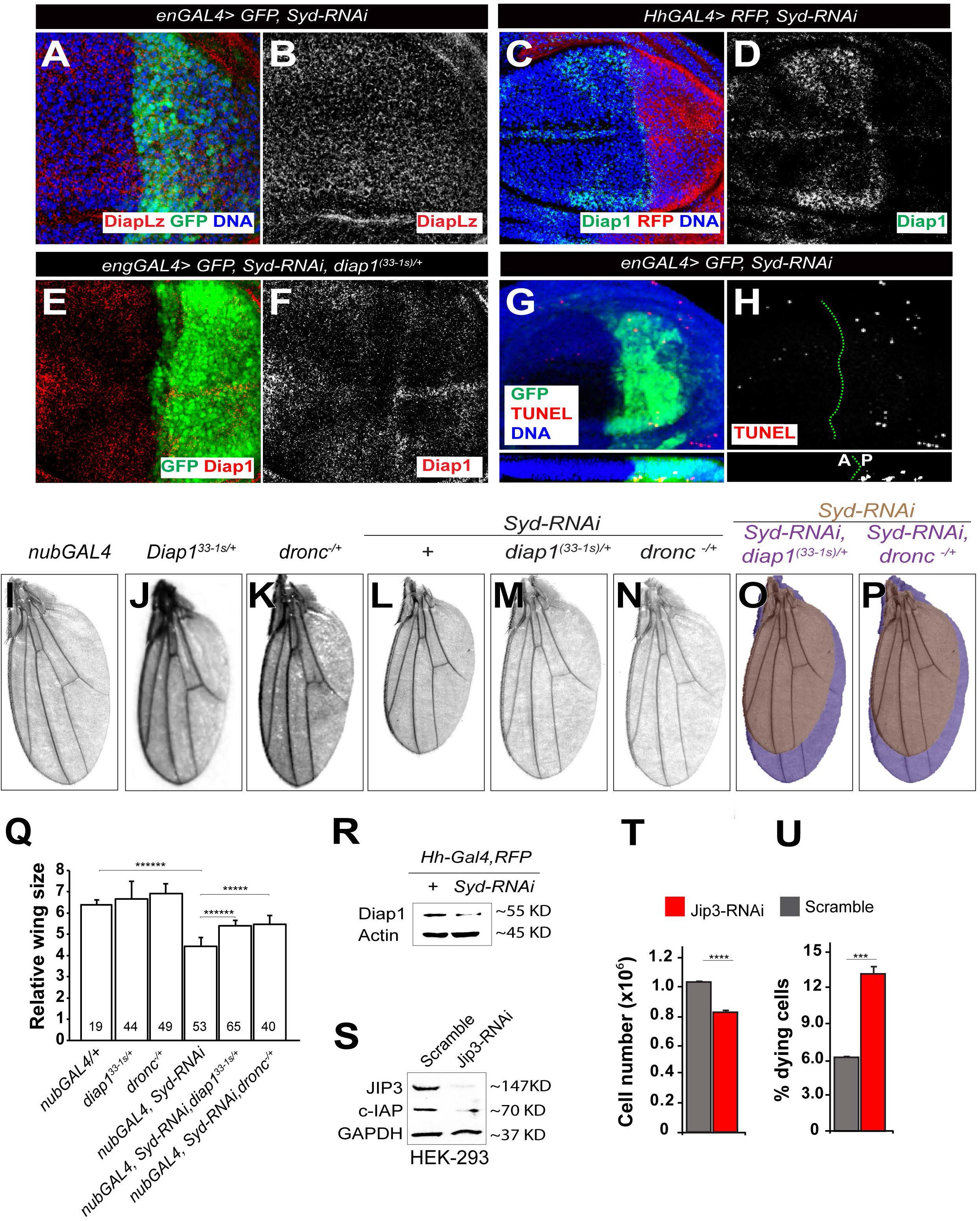
Syd/JIP3 regulates wing size by controlling Diap1 protein turnover. (A-D) Representative images of wing imaginal discs expressing *Syd-RNAi* in the posterior compartment (using *enGAL4* or *HhGAL4*) and stained with antibodies against β-gal to detect Diap1 gene activity from a genomic reporter construct, *Diap1-LacZ* (A, B) or antibodies against Diap1 to detect Diap1 protein (C, D). (E-H) Representative images of wing imaginal discs expressing *Syd-RNAi* in the posterior compartment in a *diap1^(33-1s^*) heterozygote background and stained with anti-Diap1 antibodies (E, F). Representative images of wing imaginal discs expressing *Syd-RNAi* and stained with TUNEL to detect apoptotic cells are shown in (G, H). Blue color represents DAPI stain to mark DNA. A lateral view image section is shown below (H). Green dotted lines denote anterior (A) and posterior (P) compartment boundaries. (I-P) Representative images showing adult wings from female animals of the indicated genotypes. Control wing images are shown in (I) *nubGAL4* alone, (J) *diap1^(33-1s^*^)^ heterozygotes, and (K) *dronc*^*51*^ heterozygotes. Wings derived from *nubGal4* animals expressing *Syd-RNAi* in a wild-type background or under *diap1*^*(33-1s)*^ or *dronc^51^* heterozygosity are shown in L, M, and N. The respective pseudo-color overlay images are shown in (O, P). Brown images are from (*nubGAL4, Syd-RNAi*) animals. Indigo images are from (*nubGAL4, Syd-RNAi*) animals heterozygote either for *diap1*^*(33-1s)*^ (O) or *dronc*^*51*^ (P). (Q) Quantification of (I-P) (R) Western blot images showing Diap1 protein levels in dissected third-instar wing discs expressing *srcRFP* alone (control) or co-expressing *Syd-RNAi* in the posterior compartment using *HhGAL4*. Blots were stained with anti-Diap1 or actin (loading control) antibodies. (S) Western blot images showing the effect of JIP3 knockdown on c-IAP protein levels in HEK-293T cells. Lysates were prepared from HEK-293 cells transfected either with scramble control or JIP3-siRNA. Blots were stained with anti-JIP3 antibodies to confirm Jip3 knockdown or anti-c-IAP, or GAPDH (loading control). (T) Graph showing the effect of JIP3 knockdown on HEK-293 cell numbers. (U) Graph showing the comparative proportion of propidium iodide (PI)-positive cells with or without JIP3 knockdown using flow cytometry.

We explored the possibility that Syd/JIP3 controls cellular response to Hippo signaling via a post-translational mechanism and found that it does. Although RNAi knockdown of Syd in the posterior compartment (*enGAL4, Syd-RNAi)* had no effect on Diap1 expression, immuno-staining experiments using antibodies against Diap1 protein revealed a notable reduction of Diap1 protein levels (Figure 4A-D).

Due to constraints of some genetic manipulations and for practical reasons the posterior drivers *en-GAL4* and *Hh-GAL4* are used interchangeably in the remaining experiments. We performed western blot assays using lysates prepared from wild-type larval wing tissues or tissues expressing *Syd-RNAi* in the posterior (*HhGAL4>Syd-RNAi*). Compared to wild-type controls, Syd knockdown in only half of the cells was sufficient to show a reduction of Diap1 protein levels (Figure 4R). Diap1 normally promotes cell survival by antagonizing the pro-apoptotic caspase Dronc (Nedd2-like caspase– FlyBase) (Lee et al., 2011; Wilson et al., 2002). Concordant with a loss of Diap1 protein, Syd knockdown in posterior compartment cells (*HhGAL4>Syd-RNAi*) resulted in ectopic cell death compared to anterior compartment cells within the same tissues using TUNEL assays (Figure 4G, H). Knocking down JIP3 in mammalian (HEK293) cells similarly suppressed the protein levels of c-IAP (cellular inhibitor of apoptosis), increased cell death, and reduced cell numbers (Figure 4S-U). These findings argue that Syd/JIP3 controls Diap1 turnover and that this mechanism is conserved. The E3 ubiquitin ligase function of the Diap1 RING finger domain enables Diap1 to degrade not only downstream substrates, but also itself to modulate cell survival (Ryoo et al., 2002; Yoo, 2005; Yoo et al., 2002). Similar to inhibitor of apoptosis (IAP) proteins in vertebrates, Diap1 dimerizes and self-ubiquitinates via its RING domain to trigger its own proteolysis (Feltham et al., 2011; Mace et al., 2008; Rumpf et al., 2011; Silke et al., 2005). Ectopic expression experiments have shown that inhibition of the RING domain via point mutations or deletions (RINGΔ results in a significant stabilization of not only the mutant protein but also endogenous wild-type Diap1, consistent with Diap1 dimerization (Wilson et al., 2002; Yokokura et al., 2004). RING domain defective *diap1* mutants are homozygous lethal but heterozygous viable, presumably because RINGΔwild-type Diap1 heterodimers still retain some ability to inhibit Dronc and promote cell survival (Lisi et al., 2000; Wilson et al., 2002; Yokokura et al., 2004). We asked to what extent diap1 RINGΔ heterozygosity restores Diap1 protein levels in Syd/JIP3 knocked down cells.

We took advantage of a heterozygous viable *diap1* RINGΔ mutation [*diap1^(33-1s^*^)^] (Wilson et al., 2002) and asked whether its heterozygosity rescues Diap1 in Syd/JIP3-depleted wings. Heterozygosity of the *diap1*^*(33-1s)*^ allele [*diap1^(33-1s^*^)/+^] restored Diap1 protein levels in *Syd-RNAi* expressing cells (*enGAL4>Syd-RNAi, diap1^(33-1)s/+^*, Figure 4C-F; 71%, *N=14*) and correspondingly rescued wing size (Figure 4I-M, O, Q). Thus, Syd/JIP3 controls tissues size via Diap1.

Also, given the loss of Diap1 protein in Syd/JIP3-depleted tissues and the resulting ectopic cell death, this would elevate Dronc activity. We predicted that Syd/JIP3-depleted tissues would be sensitive to Dronc dosage such that a partial reduction of Dronc function (Dronc haploinsufficiency, *dronc^51^*^/+^) will be sufficient to restore wing size to some extents. Indeed, heterozygosity of a *dronc* null allele (*dronc*^*51*^) rescued wing size in *Syd-RNAi* animals (Figure 4I, K, L, N, P, Q).

Finally, we turned to HEK-293 cells to assess whether Syd/JIP3 similarly regulates cell survival and cell growth in mammalian cells. RNAi knockdown of JIP3 inhibited the growth of HEK-293 cells and caused ectopic cell death in flow cytometric assays (Figure 4S-U), mimicking Syd/JIP3 knockdown in *Drosophila*. Collectively, these findings highlight a novel and conserved Syd/JIP3 growth control mechanism. Our data indicate that Syd/JIP3 controls tissue size downstream of the Yorkie/YAP by regulating Diap1 protein turnover, hence modulating cell survival and ultimately tissue size.

## Discussion

Our understanding of the molecular underpinnings of organ size regulation is incomplete. Genetic screens in *Drosophila* imaginal discs have led to the identification of Hippo signaling as a central and evolutionarily conserved growth control mechanism. More recently Willsey and colleagues reported that in *Drosophila*, developmentally encoded signals trigger JNK activation at the posterior-anterior boundary of the developing wing. This localized JNK activity, broadly inhibits Hippo signaling to promote tissue-wide cell proliferation, resulting in a properly sized adult wing (Willsey et al., 2016). Here we show that Syd/JIP3 regulates wing size downstream of Yorkie by controlling Diap1 protein turnover.

Direct activation of Yorkie/YAP potently triggers tissue overgrowth in vertebrates and *Drosophila*. Syd knockdown partly suppresses Yorkie/YAP-mediated tissue overgrowth. The moderate effect is likely because direct activation of Yorkie/YAP is so potent that the resulting ectopic cell proliferation masks the impact of *Syd-RNAi/Diap1*-mediated cell death in these tissues. Consistent with this, depletion of Syd protein under three independent experimental conditions (*Syd-RNAi* or Syd haploinsufficiency using *syd*^*PL00426*^ or *Syd^A2^* loss of function alleles) suppressed the large wing phenotype caused by a more moderate Yorkie activation (Hippo knockdown).

Activation of Yorkie/YAP either via Hippo knockdown or direct Yorkie/YAP overexpression resulted in the upregulation of Syd/JIP3 protein in Drosophila and mammalian cells. These findings are consistent with Syd/JIP3 acting downstream of Yorkie/YAP signaling.

Syd/JIP3 regulates growth by controlling Diap1 protein stability. Inhibition of Syd/JIP3 decreases Diap1 protein levels and correspondingly triggers ectopic cell death in *Drosophila* and in mammalian cells. Similar to *Syd-RNAi*, expression of *Diap1-RNAi* in P/A cells is sufficient to reduce adult wing size (Supplemental Figure 6). Partial restoration of Diap1 rescues the small wing effect of Syd/JIP3 inhibition. The partial rescue may reflect an incomplete restoration of Diap1 protein and/or that the re-established pool of Diap1 is only partly effective at inhibiting cell death. We cannot rule out the possibility that Syd/JIP3 tissue size regulation involves additional Diap1-independent mechanisms. The observation that Diap1 RINGΔ heterozygosity restores Diap1 protein levels in Syd/JIP3 knockdown cells supports the notion that Syd/JIP3 normally stabilizes Diap1 by protecting it from proteolytic degradation.

Ectopic cell death is the primary mechanism underlying the reduced growth of Syd knockdown *Drosophila* tissues and human cells. We did not detect a decrease of proliferative capacity in *Syd-RNAi* expressing *Drosophila* tissues compared to wild-type controls (Supplemental Figure 5).

This study has identified Syd/JIP3 as a novel size regulator downstream of Hippo. It is tempting to speculate that Syd/JIP3 regulation of Diap1 likely represents a robustness mechanism for Yorkie-mediated tissue growth. *Syd/JIP3* is expressed at higher levels during the rapid growth phase (72-96 hours after egg laying-AEL) and progressively drops later (120-150 hours AEL) when growth slows down (Supplemental Figure 7). High Syd/JIP3 levels would stabilize Yorkie-activated Diap1, permitting sustained tissue growth phase. The decline of Syd/JIP3 transcription later in development would target Diap1 for degradation, thereby allowing cell death to sporadically occur throughout the tissue, hence scaling down tissue response to Yorkie and contributing to size stereotypy.

Additional significance can be derived from the fact that our findings provide potential mechanistic insights into the recent and perplexing link between *JIP3* mutations and organ size defects in mammals, including humans where de novo *JIP3* variants are associated with microcephaly (Kelkar et al., 2003; Platzer et al., 2019).

## Acknowledgment

We thank A. Bergmann, G Halder, T. Xu, H. Steller, M. Baylies and D. Pan for antibodies and fly stocks. We also thank BDSC and VDRC for fly lines. We also thank M. Kroll for comments on the manuscript.

## Author Contributions

V.A and C.Y.C designed the research. V.A performed *Drosophila* experiments and analyzed the data together with C.Y.C. G.P.V performed tissue culture experiments. C.Y.C wrote the manuscript.

---

**Supplemental Figure 1.**
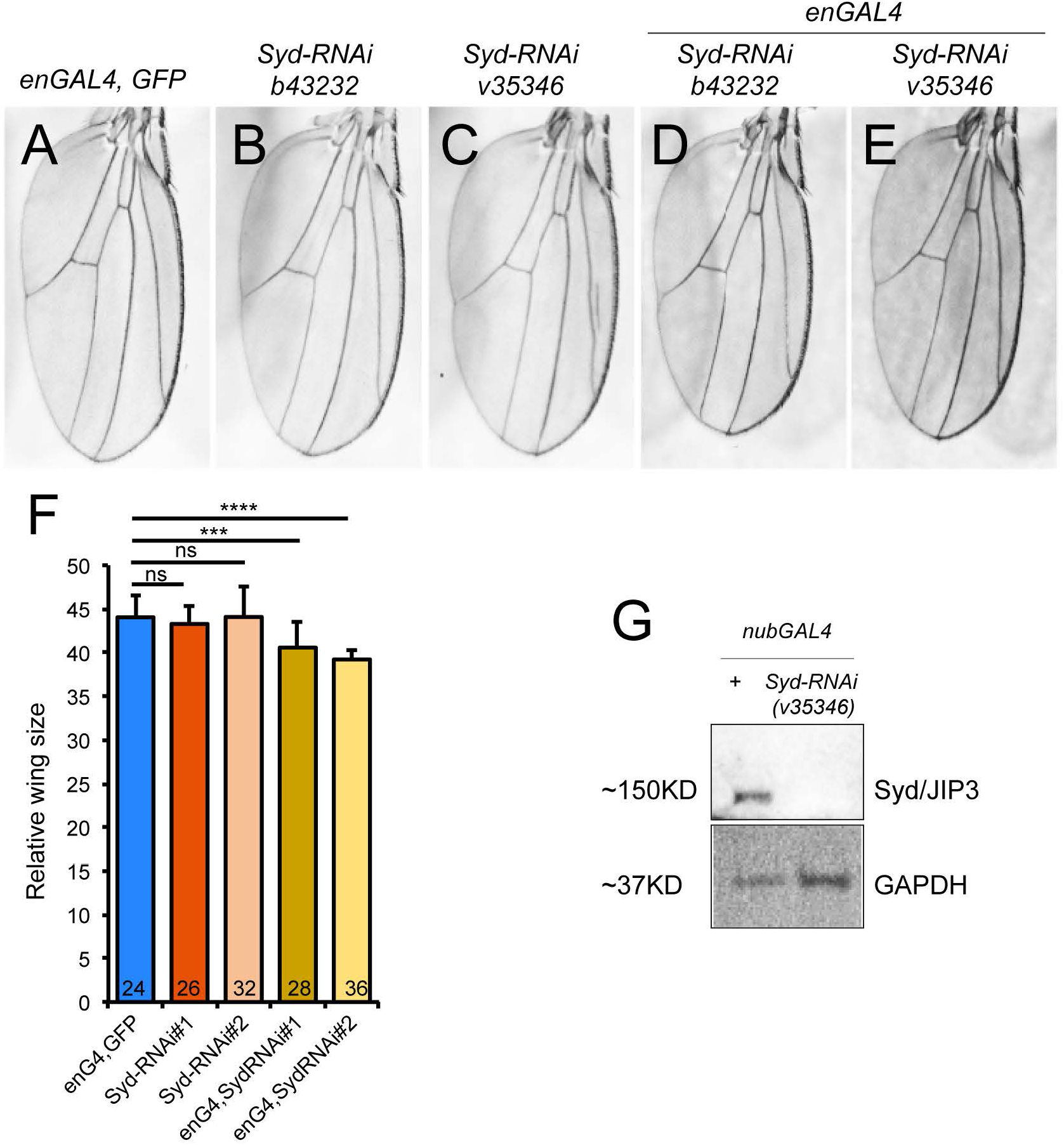
Syd/JIP3 knockdown in the posterior compartment suppresses adult wing size. (A-F) Representative images showing adult wings from female animals of the indicated genotypes (A-E). Syd knockdown effect on wing size using *Syd-RNAi* (*b43232*) (A, B, D, F) or *Syd-RNAi* (*v35346*) (A, C, E, F). (F) Quantification of (A-E).

**Supplemental Figure 2.**
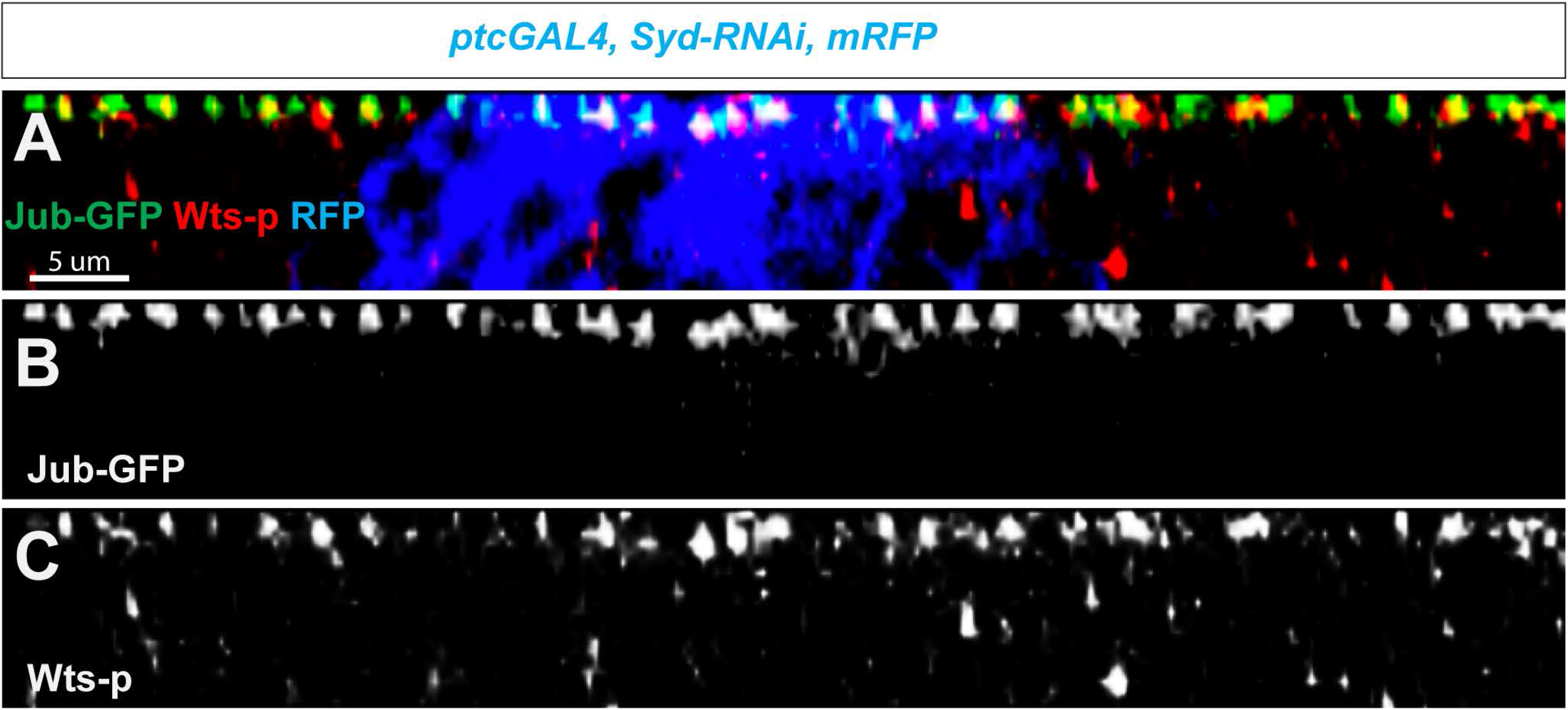
Expression and subcellular localization of ajuba and Warts in Syd knockdown cells. (A-C) Lateral section images showing wing imaginal discs containing a genomic rescue construct expressing Ajuba tagged with GFP under its own promoter (*Jub-GFP) (Sabino et al., 2011)* and co-expressing *Syd-RNAi* and RFP (shown in blue) under *ptcGAL4.* Tissues were stained with anti-phosphorylated Warts (pWts) antibodies.

**Supplemental Figure 3.**
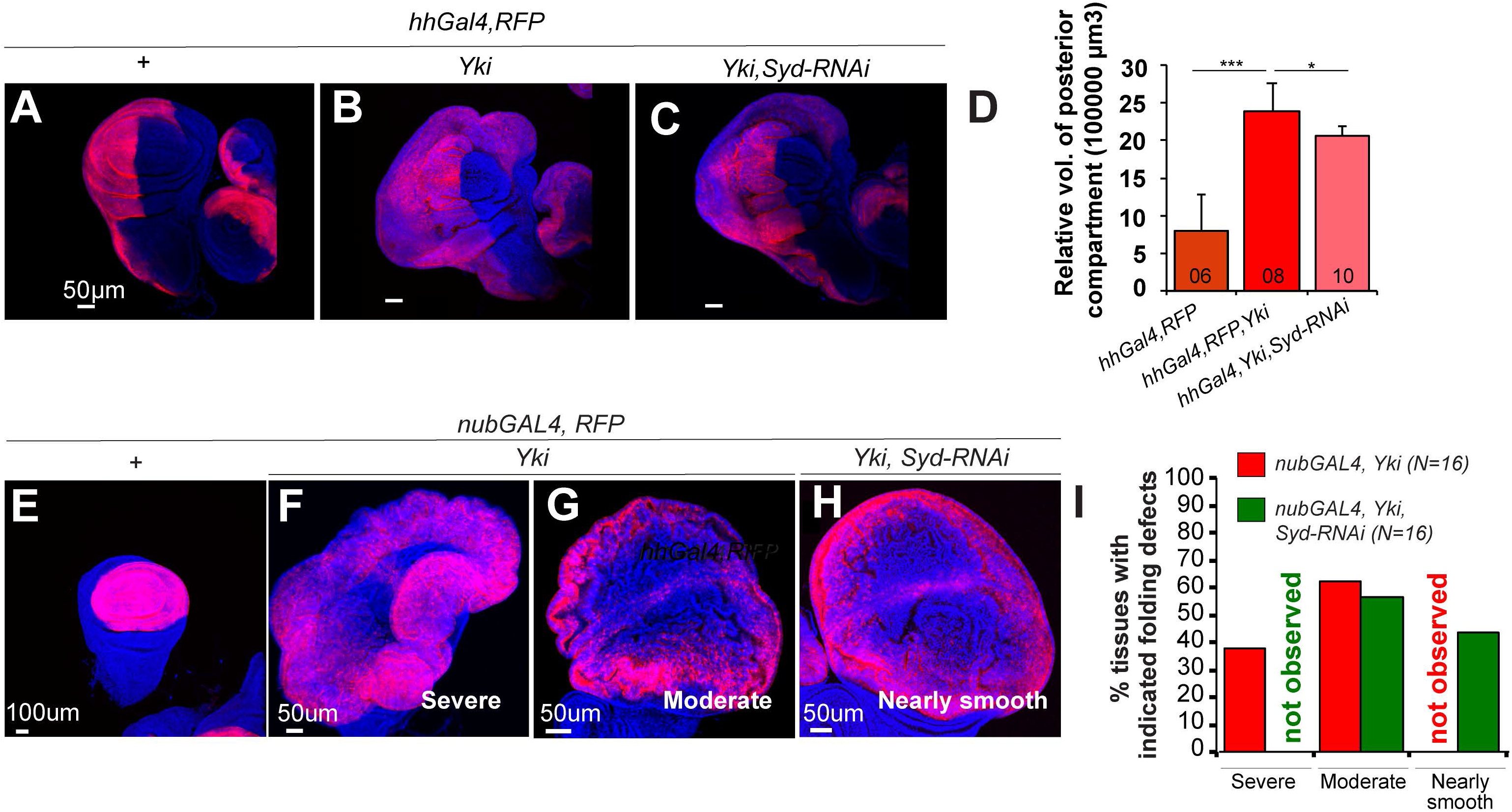
*Syd/JIP3* knockdown suppresses *Yorkie*-mediated tissue overgrowth. (A-D) Confocal images of wing imaginal discs from age-controlled animals expression *RFP* (A) alone or co-expressing *Yorkie* (B) or co-expressing *Yorkie* and *Syd-RNAi* (C) under the control of posterior driver *HhGAL4*. (D) Volume quantification of *RFP*-positive tissues in (A-C). Yorkie overexpression caused overgrowth in the posterior compartment and showed an increased RFP-positive area, RFP intensity and tissue folding. Syd depletion (*HhGAL4,Yorkie, Syd-RNAi*) reduced the size of RFP-positive tissues. (E-I) Images of wing imaginal discs from age-controlled animals expressing *RFP* alone (E) or co-expressing *Yorkie* (F and G) or co-expressing *Yorkie* and *Syd-RNA I* (H) under the control of *nubGAL4*. (I) Quantification of (E-H). *Yorkie* overexpression results in overgrown tissues showing severe or moderate tissue folds, 37% and 63% (N=16), respectively. Syd knockdown (*nubGAL4, Yorkie, Syd-RNAi*) suppressed the appearance of tissue folds and yielded tissues with moderate to smooth folding (*N* =16, 56% and 44%, respectively). Severe tissue folding was not detected in (*nubGAL4, Yorkie, Syd-RNAi*) tissues (H).

**Supplemental Figure 4.**
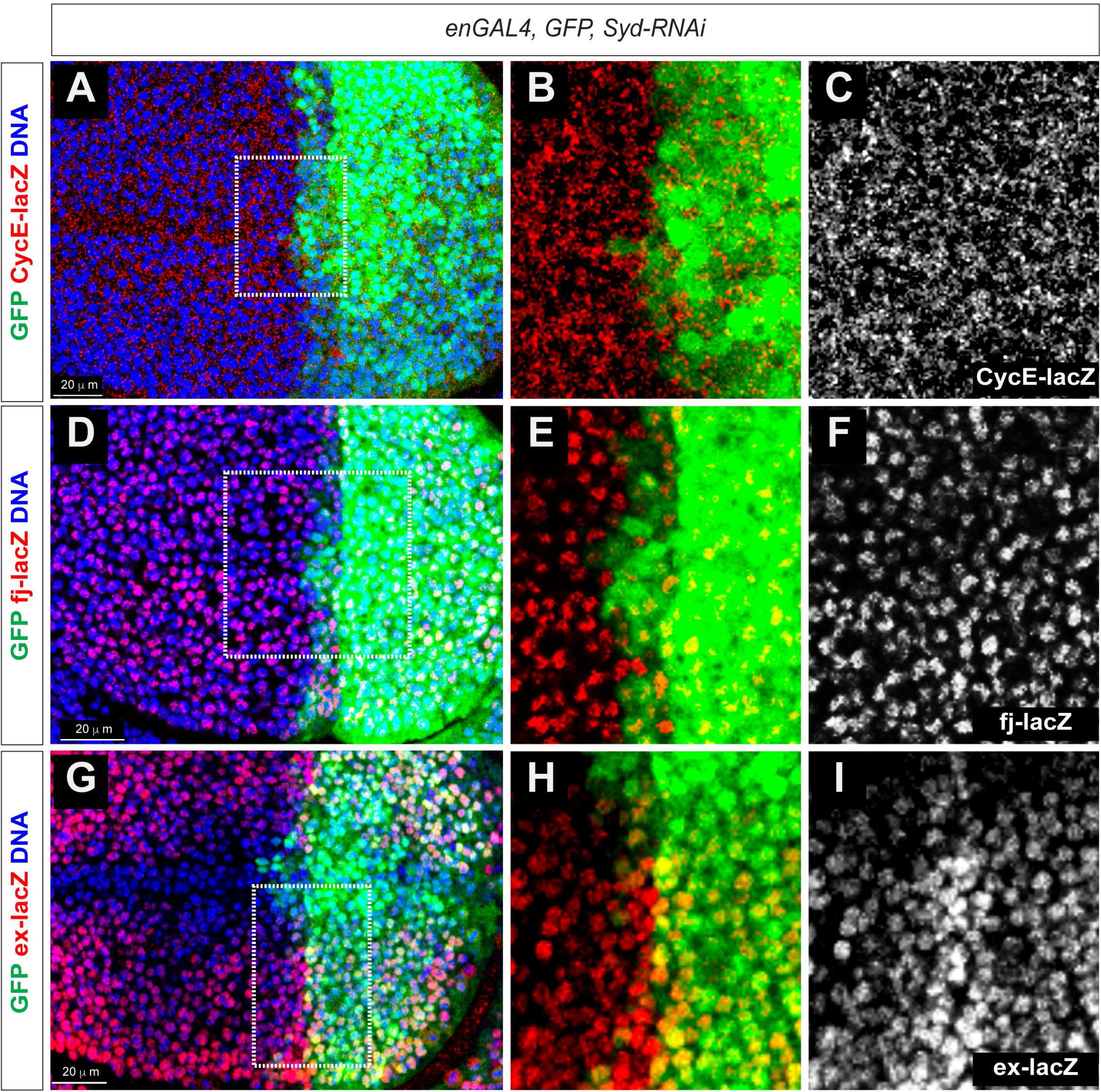
Transcription of *four-jointed* and *cyclin-E* in Syd knockdown cells. (A-I) Representative images of wing imaginal discs dissected from *four-jointed-LacZ* or *CyclinE-LacZ* or *Expanded-LacZ* reporter lines expressing *Syd-RNAi* in the posterior compartment using *enGAL4* and stained with antibodies against β-gal to monitor gene activity of *cyclinE* (A-C) or *four-jointed* (D-F) or *expanded* (G-I). The insets show that *cyclinE* (B, C) is localized in nucleus while *four-jointed* (E, F) and *expanded* (H, I) are localized at membranes.

**Supplemental Figure 5.**
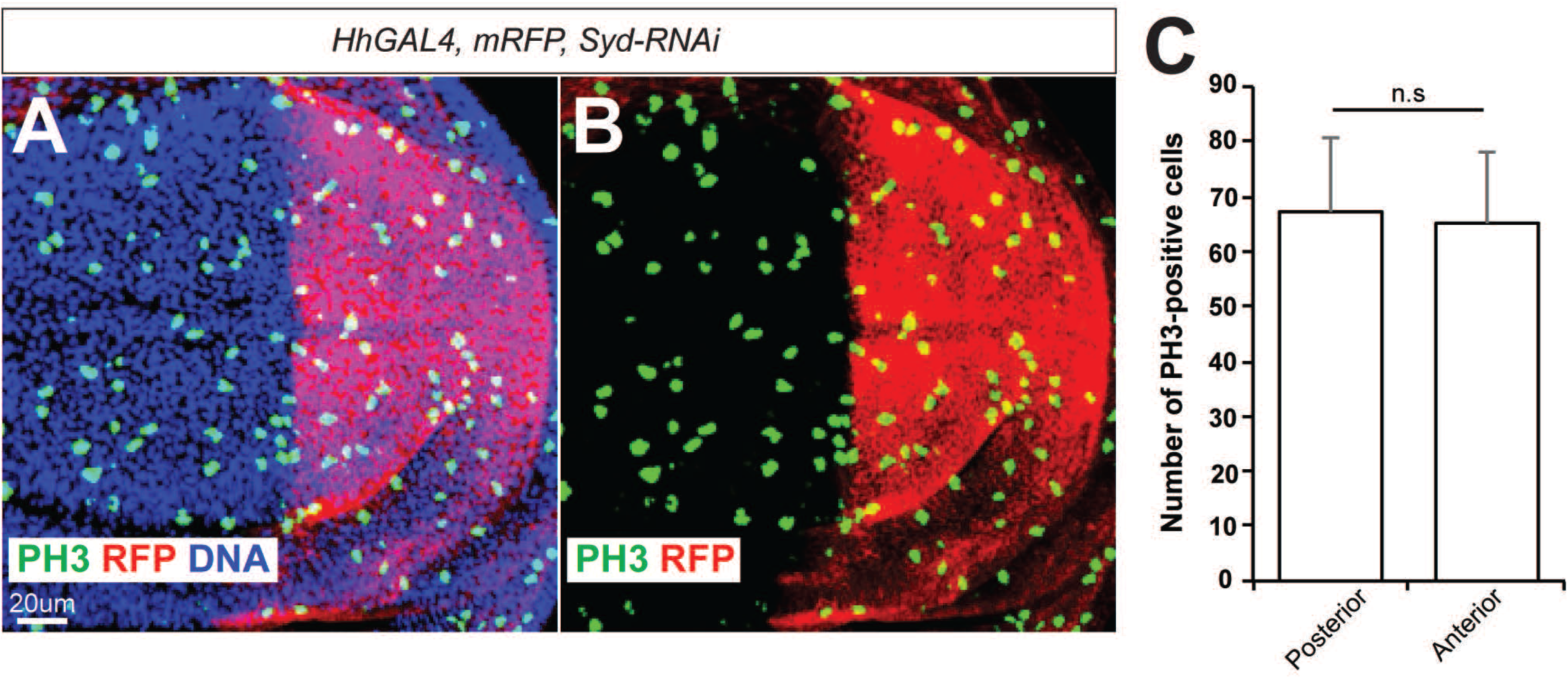
Mitotic potential of Syd depleted cells. (A, B) Image showing a wing imaginal disc expressing *Syd-RNAi* under the *HhGAL4* driver and stained with anti-phosphorylated Histone-3 to detect proliferating cells. (C) Quantitation of (A, B). *P*-value (t-test) was 0.78.

**Supplemental Figure 6.**
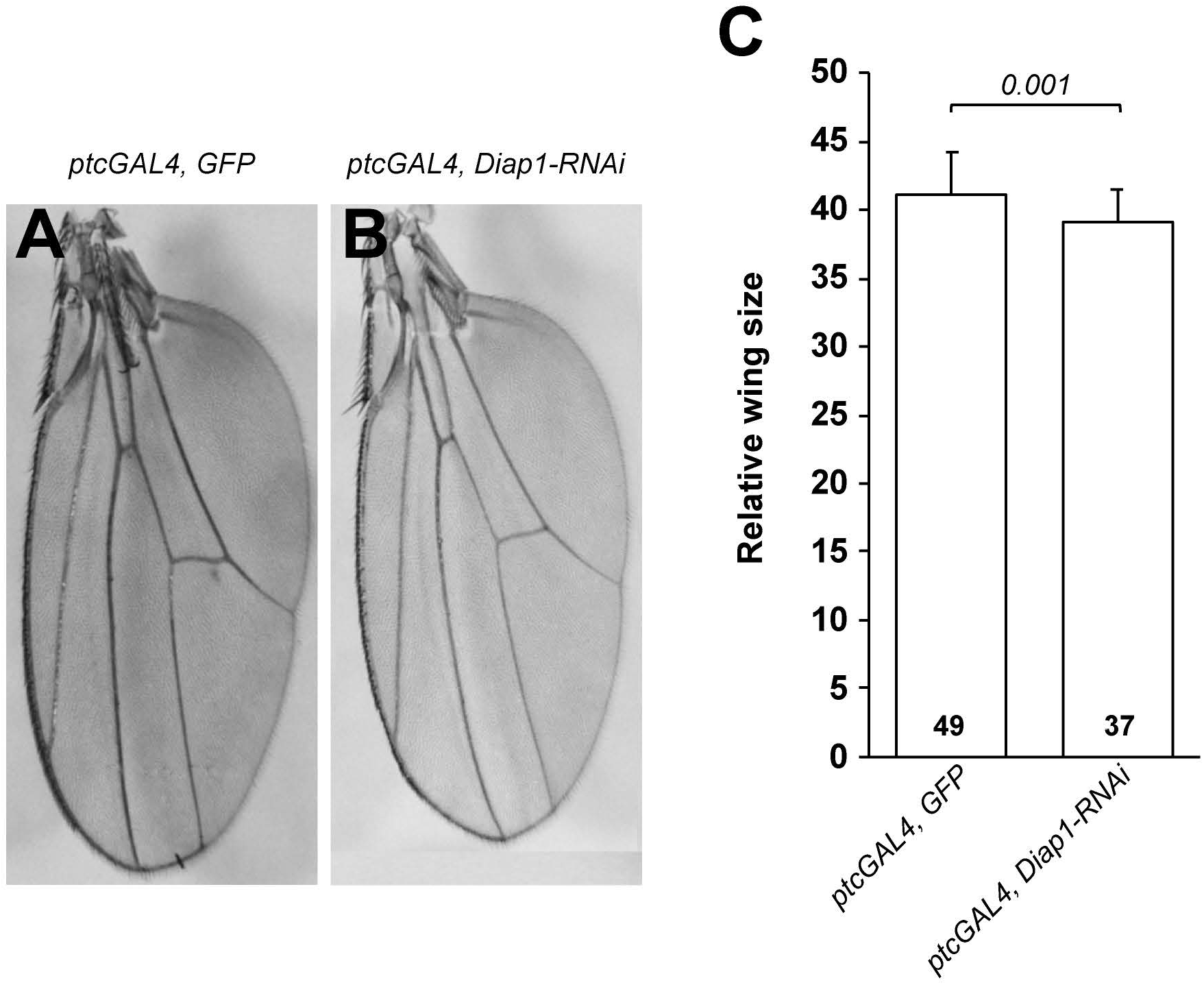
Effect of Diap1 knockdown on adult wing size. (A, B) Images showing wing imaginal discs dissected from female animals of the indicated genotypes. (C) Quantification of (A, B).

**Supplemental Figure 7.**
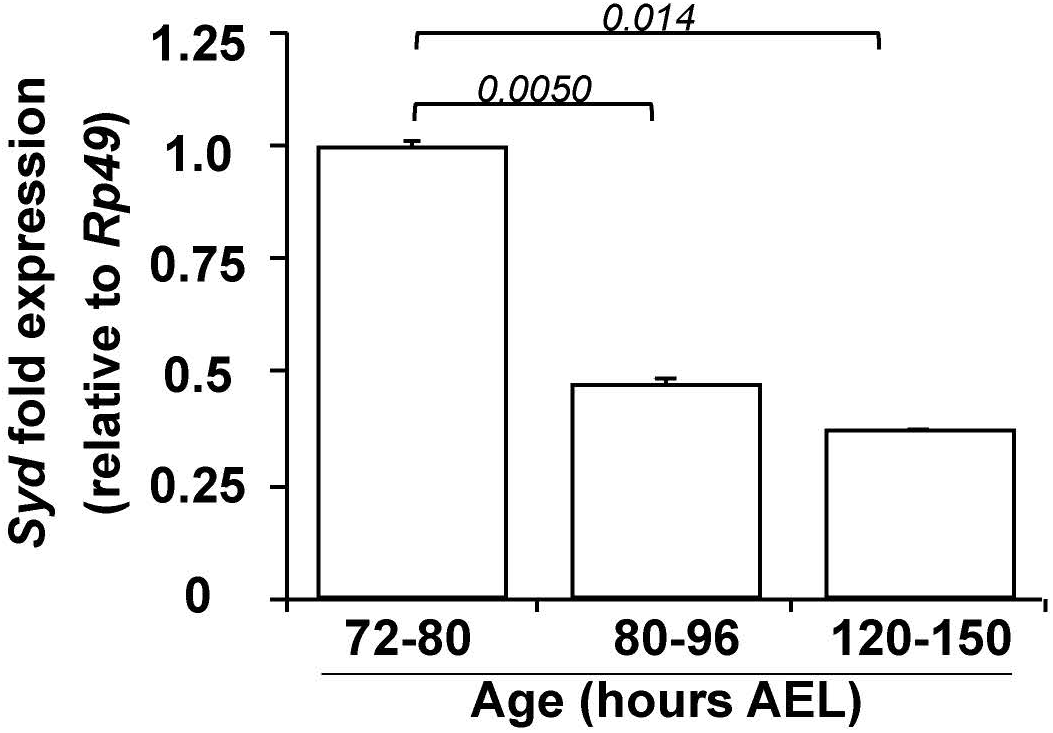
*Syd* gene expression during wing development. Graph showing *Syd/JIP3* expression in wing imaginal discs dissected at the indicated developmental time periods from biological triplicates. *Syd/JIP3* expression was calibrated to *Rp49* expression at each developmental time period. Fold expression changes were established by comparing each time period to the 72-80 hours AEL period. Graph shows mean fold changes in gene expression ± SD.

**Supplemental Figure 8.**
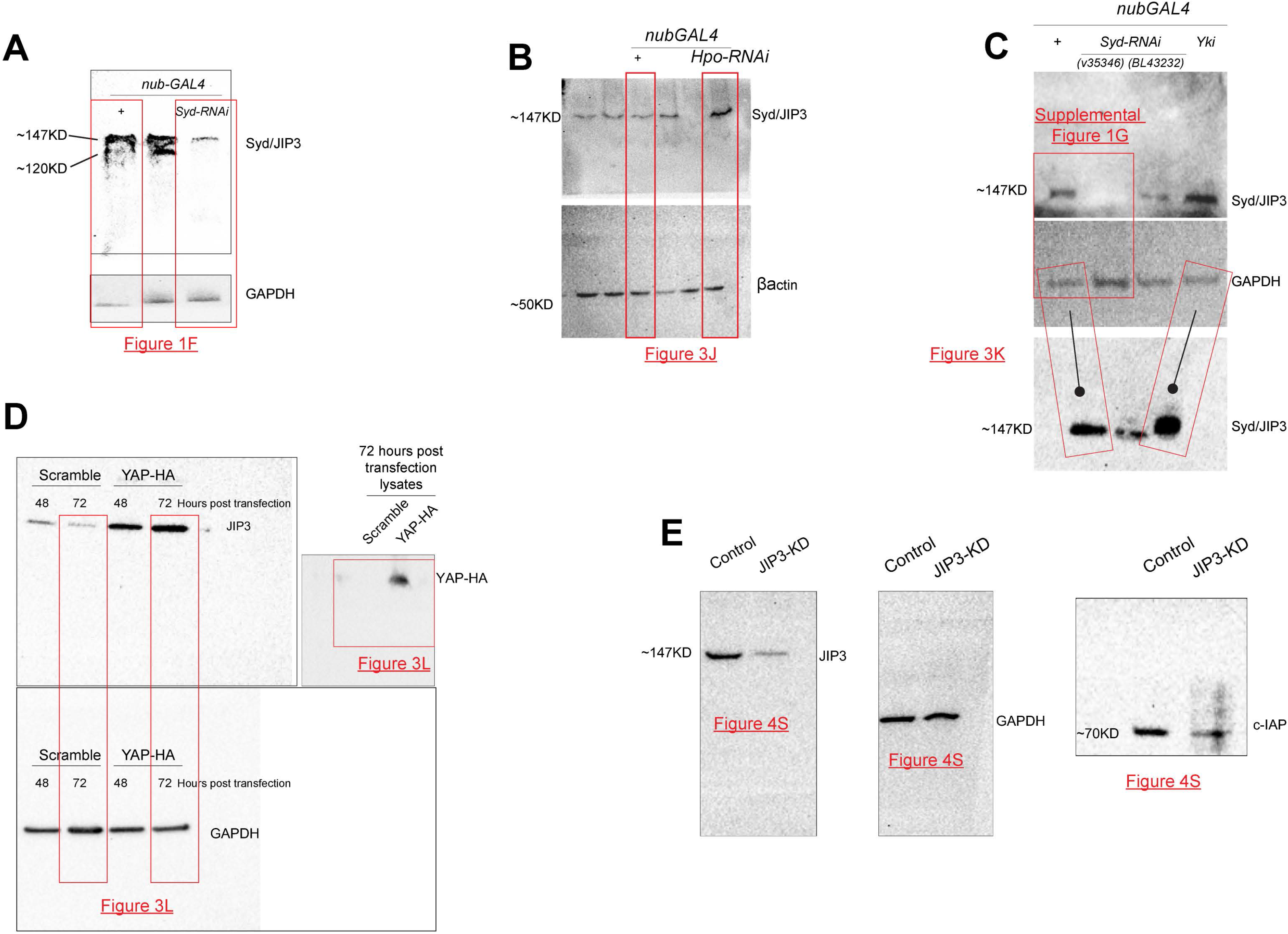
Full image of presented Western blots. Boxes denotes the data presented in the indicated figures (underlined red text) (A) Full Western blots related to Figure 1F. Anti-JIP3 Western blots from *nubGAL4* (control) or Syd-knockdown (*nubGAL4, Syd-RNAi*) wing tissue lysates. Anti-GAPDH was used as an input control. In *Drosophila*, anti-human JIP3 antibody detected two bands: a prominent ~147KD band and a much weaker 135KD band in lysates derived from wild-type wing tissues. Both of these bands were considerably reduced in lysates obtained from wing tissues expressing Syd-RNAi. Note that the sparse 135KD JIP3 band was not always detected in lysates with lower protein yield. (B) Related to figure 3J. Uncropped images of anti-Syd/JIP3 Western blotting using controls (*nubGal4*) or Hippo knockdown (*nubGal4, HpoRNAi*) larval wing tissue lysates. β-actin is a loading control. (C) Related to figure 3K. Uncropped anti-Syd/JIP3 Western blot image showing Syd/JIP3 protein levels in control (lane#1, *nub-Gal4*) tissues or tissues expressing *Syd-RNAi* (lanes 2 and 3) or overexpressing Yorkie (*nubGal4,Yorkie*) (lane 4). GAPDH is a loading control. (D) Related to figure 3L. Uncropped Western blot images of showing upregulation of JIP3in HEK293 cells overexpressing YAP (YAP-HA) compared to scramble controls. GAPDH is a loading control. (E) Related to figure 4S. Uncropped Western blot images showing the effect of JIP3 knockdown on c-IAP expression in HEK 293 cells.

